# Alfalfa intercropping maintains intermediate wheatgrass grain yield relative to nitrogen fertilization across site conditions and stand ages

**DOI:** 10.64898/2026.10.21.759482

**Authors:** Kenneth C. Okonkwo, Thomas P. McKenna, Molly Hanlon, Shannon Meehan, Nia Worth, Tyler Thrash, Timothy E. Crews, Allison J Miller

## Abstract

Intermediate wheatgrass (IWG; *Thinopyrum intermedium*) is a promising perennial grain crop, but yield instability remains a major challenge. We compared effects of nitrogen fertilization and intercropping with alfalfa (*Medicago sativa* L.) on IWG biomass, grain yield, and root traits across three stand ages at two nearby sites in eastern Missouri differing in soil properties (Planthaven Farms and Danforth Center Field Research Site). Four treatments (unfertilized IWG, fertilized IWG, IWG-alfalfa intercrop, and alfalfa monoculture) were established in a randomized complete block design with four replications per site. Biomass and grain yield were evaluated at per-row and field scales, while roots were measured to 1 m depth during the second stand age. Alfalfa intercropping maintained IWG grain yield without synthetic N fertilizer, although treatment responses varied by site and stand age. At Planthaven, differences became more pronounced as stands matured, with intercropped IWG producing the greatest per-row biomass and grain yield in stand age three. At the Danforth Center Field Research Site, intercropped IWG produced greater per-row biomass and grain yield than monocultures, whereas fertilized IWG tended to produce greater field-scale biomass; grain yield declined across treatments with stand age. Across sites, nitrogen fertilization increased vegetative biomass more consistently than grain yield. Despite having fewer IWG rows, intercrops maintained field-scale grain yield comparable to fertilized IWG in several site-years. Root traits responded weakly to management and were structured primarily by soil depth. These findings highlight the importance of site conditions, and stand age, when evaluating management strategies for perennial grain production.

## 1. INTRODUCTION

Perennial grains are receiving increasing attention as concerns mount about impacts of agriculture on soil (Lal, 2015; Bogužas et al., 2022). Through their deep, persistent root systems and continuous living cover, perennials offer reduced soil disturbance, erosion, and nitrate leaching, and contribute positively to soil carbon storage (Crews et al., 2018; van der Pol et al., 2022). Intermediate wheatgrass (IWG; *Thinopyrum intermedium* (Host) Barkworth & D.R. Dewey; trade name Kernza^(R)^) is a leading perennial grain candidate under active domestication for both edible grain and forage production (Bajgain et al., 2022). IWG breeding has improved grain size, shattering resistance, threshability, harvestability, and grain yield (Wagoner, 1990; DeHaan et al., 2018; Crain et al., 2024). Nevertheless, IWG grain yields remain below those of annual cereals and often decline after the first production year (Jungers et al., 2017; Fernandez et al., 2020). Relatively low yields likely result from a combination of genetic, physiological, and agronomic factors, including altered resource allocation patterns (Hunter et al., 2020; Zhen et al., 2024), changes in tillering (Fernandez et al., 2020), and progressive nitrogen limitation with stand age (Jungers et al., 2017), among others.

Agronomic strategies such as nitrogen fertilization and legume intercropping have been investigated to improve IWG yield. Fertilization can increase biomass and grain yield, particularly during early production years, but responses vary with stand age and environment (Jungers et al., 2017; Shoenberger et al., 2025). However, nitrogen fertilization may also increase lodging susceptibility, potentially reducing harvest efficiency and grain recovery (Jungers et al., 2017; Frahm et al., 2018; Fernandez et al., 2020).

Intercropping IWG with legumes such as alfalfa (*Medicago sativa* L.) may provide biologically fixed nitrogen and support yield as stands mature, although outcomes depend on intercrop design (addition or replacement designs) and environmental conditions. In addition,designs, the original IWG row spacing and density are maintained, with alfalfa planted between existing IWG rows; in replacement designs, every other IWG row is replaced with alfalfa, resulting in half the number of IWG rows. Previous replacement-design studies have reported lower IWG grain yield in the intercrop than monocultures during early stand age, followed by improved intercrop performance relative to monocultures as stands matured. Mårtensson et al. (2022) found that the IWG partial land-equivalent ratio exceeded 1 after the first production year, while Tautges et al. (2018) reported that intercrop grain yield became similar to or greater than the fertilized monoculture at several sites by the fourth production year. Other studies have reported greater biomass or improved yield stability in the IWG-alfalfa systems relative to fertilized monocultures as stands mature (Crews et al., 2022; Pinto et al., 2024). These studies indicate that IWG responses to alfalfa intercropping can change with stand age.

Differences in responses may reflect local environmental conditions, which can interact with management approaches to further influence IWG yield (Jungers et al., 2017; Pinto et al., 2024). Multi-site studies have documented substantial variation in IWG biomass and grain yield associated with soil and climate conditions, often with smaller or inconsistent responses to fertilization (Cassani et al., 2025; McKenna et al., 2026). Vapor pressure deficit has also been associated with variation in IWG yield across space and time (Zhen et al., 2024). Although McKenna et al. (2026) demonstrated variation in responses to nitrogen fertilization and alfalfa intercropping across geographically diverse sites, less is known about whether similar variation occurs between nearby sites within the same climatic region. Understanding this finer-scale variation is important for determining whether management recommendations can be transferred among fields within the same region, but with contrasting local conditions.

Additionally, belowground traits may provide insight into IWG responses to intercropping and fertilization, yet roots are rarely measured alongside aboveground yield. Root length, root length density, diameter, and biomass influence water and nutrient acquisition and competitive interactions in mixed stands (Ravenek et al., 2016; Lynch, 2022). In perennial grasses, nitrogen fertilization often shifts allocation toward shallower rooting, while intercropping can alter root distributions through niche differentiation, potentially changing where and how plants acquire resources (Houde et al., 2020; Homulle et al., 2022).

In IWG, nitrogen deficiency can alter root allocation and increase root-to-shoot ratio under controlled conditions (Dobbratz et al., 2023; Brekalo et al., 2024), whereas field fertilization can alter root N concentration and nitrogen-use efficiency without significantly changing root biomass (Sprunger et al., 2018; Dobbratz et al., 2023). IWG root biomass and N-use patterns also vary with stand age and season, and relationships between root traits and aboveground yield can be weak (Griffin et al., 2024). In IWG-alfalfa intercrops, Crews et al. (2022) found greater total root biomass in the upper 10 cm of IWG-alfalfa intercrops than in fertilized or unfertilized IWG monocultures, and Li et al. (2019) reported that root biomass responses in varied with N availability and species composition under controlled conditions. However, evidence for how legume intercropping affects IWG root traits, and spatial distribution remains scarce, particularly alongside yield responses across field environments.

These knowledge gaps limit our understanding of how local environmental conditions and stand age shape above- and below-ground responses of IWG to nitrogen fertilization and legume intercropping. In particular, it remains unclear whether responses diverge between nearby sites within the same climatic region and whether differences in aboveground responses are accompanied by changes in root traits.

In this study, we examined two nearby eastern Missouri sites within a seven-site experiment (McKenna et al., 2026), incorporating additional stand years and root measurements. Specifically, we assessed (i) biomass and grain yield responses to fertilization and intercropping across stand ages and measurement scales; (ii) root trait responses across treatments, soil depths, and sampling positions; and (iii) differences in treatment responses between sites. We hypothesized that (i) both nitrogen fertilization and alfalfa intercropping would increase IWG biomass and grain yield, but their effects would vary with stand age, with the relative benefit of intercropping increasing as stands matured; (ii) fertilization and intercropping would alter root traits and their spatial distribution; and (iii) treatment responses would differ between sites.

## 2. MATERIALS and METHODS

### 2.1 Study Sites

The study was conducted at two sites in eastern Missouri, Planthaven Farms in O’Fallon, MO and the Danforth Center Field Research Site (DDPSC FRS) in St. Charles, MO; both sites are part of a larger seven-site experiment (McKenna et al., 2026). The two sites used in this study are 31 km apart and experience cold winters and warm, humid summers. Both were previously managed under conventional tilled corn-soybean rotations. The IWG-alfalfa experiment was established at Planthaven Farms in September 2021. In August 2022, the Danforth Center acquired the DDPSC FRS, and shortly thereafter in September 2022 the experiment planted at Planthaven Farms was replicated at the DDPSC FRS.

Planthaven Farms has high-clay, seasonally saturated soils classified as fine, smectitic, mesic aeric chromic vertic epiaqualfs, whereas DDPSC FRS has well-drained soils classified as coarse-silty, mixed, superactive, mesic cumulic hapludolls (USDA-NRCS, 2023). Mean annual precipitation ranges from 889 to 1195 mm, and mean annual temperature from 10.0 to 13.9°C across the sites (Table 1). Daily precipitation and mean temperature data for each site were obtained from the PRISM Climate Group, Oregon State University, using site coordinates for 2021-2025. Daily values were aggregated into monthly and annual precipitation totals and mean temperatures to characterize climatic conditions during establishment and stand development (“PRISM Group at Oregon State University,”).

**Table 1.**
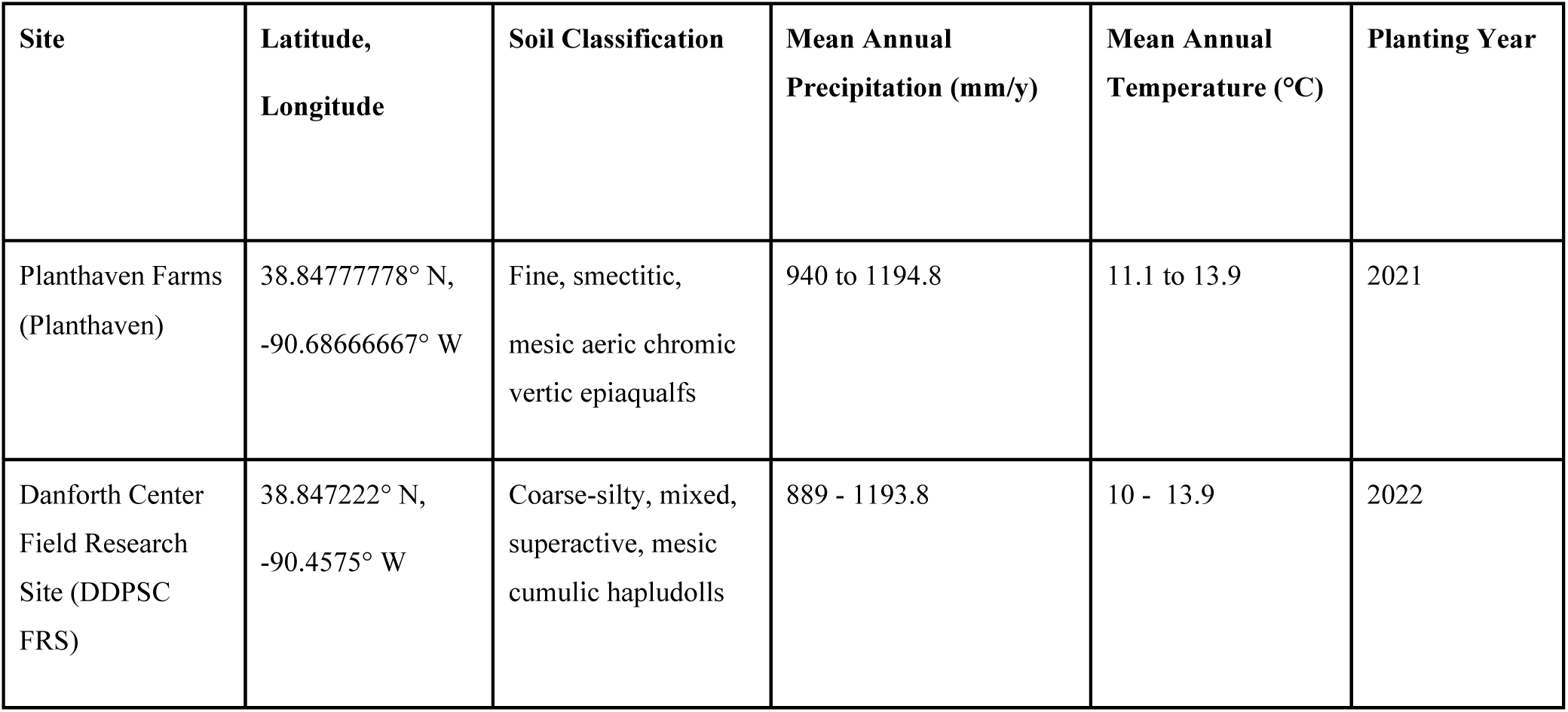
Site characteristics for the two experimental locations.

### 2.2 Experimental Design

At each site, the experiment followed a randomized complete block design with four blocks and four randomly assigned treatments: (i) unfertilized IWG monoculture (IWG), (ii) nitrogen-fertilized IWG monoculture (IWG+N), (iii) unfertilized IWG+alfalfa intercrop (IWG+Alf), and (iv) unfertilized alfalfa monoculture (Alf). Each plot was 6.096 m long and contained 18 rows spaced 0.3078 m apart. The intercrop followed a replacement design, with nine IWG rows alternating with nine alfalfa rows, with the same row spacing and total number of rows as the monocultures (Rodriguez et al., 2020). Plots were seeded in September 2021 at Planthaven Farms and September 2022 at DDPSC FRS using IWG line TLI 801 (The Land Institute, Salina, Kansas) and alfalfa cultivar Hybriforce 3400 (Dairyland Seed, USA). Both species were seeded at 15 lb pure live seed acre⁻¹. Urea was applied annually to IWG+N plots at 90 kg N ha⁻¹ each spring. Plots were hand-weeded throughout the growing season.

### 2.3 Sample Collection, Processing, and Field Observations

Before establishment, soil samples were collected from each plot to a depth of 25 cm. Samples were analyzed by the Kansas State University Soil Testing Laboratory (Manhattan, KS, USA) for soil pH, Sikora buffer pH, cation exchange capacity (CEC), potassium, calcium, sodium, organic matter (loss-on-ignition), Bray-extractable phosphorus, total nitrogen, total phosphorus, and total organic carbon.

IWG grain and aboveground biomass were harvested annually at full grain maturity, typically in late July or early August, for three production years (2022-2024 at Planthaven Farms; 2023-2025 at DDPSC FRS). Seed heads were clipped from 3m sections of three representative IWG rows per plot (9 linear m total), followed by harvest of the remaining aboveground vegetative biomass from the same sections. Seed heads were excluded from vegetative biomass measurements. The harvested vegetative biomass was weighed in the field, and subsamples were oven-dried at 60°C for 48 h to determine dry mass. IWG Seed heads were similarly dried and threshed at The Land Institute (Salina, KS, USA) using a Wintersteiger LD-50 laboratory thresher. Threshed grain was weighed to determine grain yield. Biomass and grain yield were quantified at two scales: per-row yield (g m⁻²), representing yield of the sampled IWG rows, and field-scale yield (kg ha⁻¹), accounting for IWG row density. For per-row yield, harvested mass was divided by the sampled row length (9 m) and row spacing (0.3048 m). Field-scale yield was calculated by multiplying per-row yield by 10 for monocultures and by 0.5 for the intercrop, where IWG occupied half the rows. IWG lodging was visually assessed as the percentage of the plot canopy within three stem-angle classes relative to the soil surface: 70-45°, 45-10°, and <10°. Lodging was assessed immediately before harvest in 2024 at both sites (stand age 3 at Planthaven Farms and age 2 at DDPSC FRS) and in May, July, and immediately before harvest in August 2025 at DDPSC FRS (stand age 3).

Alfalfa aboveground biomass was harvested when approximately 10-15% of alfalfa stems were flowering, typically three times annually, except in 2024, when severe alfalfa weevil (*Hypera postica*) infestation at both sites limited harvests to two in that year. Alfalfa was cut to a height of 10 cm along 3m sections of representative interior rows. All harvested alfalfa aboveground material, including stems, leaves and flowers, was included in biomass measurements. Samples were weighed in the field, and then subsamples were dried at 60°C for 48 h and subsequently weighed to determine dry mass. After measurement, all harvested alfalfa aboveground biomass, including subsamples, was returned to its respective plot. Repeated harvests were intended to promote nitrogen availability to IWG through root exudation, turnover, and residue return (Crews et al., 2016, 2022).

Roots were sampled in August of the second stand age at each site (2023 at Planthaven Farms; 2024 at DDPSC FRS). Four soil cores (4.45 cm diameter x 1 m depth) were collected per plot, with two within plant rows and two between rows (64 cores per site). Cores were collected using a Geoprobe at Planthaven Farms and a hydraulic Giddings probe at DDPSC FRS, then divided into three depths: 0-20, 20-50, and 50-100 cm. Roots were recovered using a GVF Hydropneumatic Elutriation System 13000, scanned at 600 dpi with an Epson 13000XL flatbed scanner, and analyzed in RhizoVision using default segmentation and analysis settings (Seethepalli et al., 2021). Roots were then dried at 60°C for 48 h and weighed to determine dry root mass. Root length density (RLD) was calculated as TRL divided by soil core volume. Specific root length (SRL) was calculated as TRL divided by root dry mass.

### 2.4 Statistical Analysis

All statistical analyses were conducted in R version 4.5.2. Sites were analyzed separately because they were established in different years. IWG vegetative biomass, grain yield, and alfalfa aboveground biomass were analyzed using separate linear mixed-effects models, with treatment, categorical stand age, and their interaction as fixed effects and block as a random intercept. IWG biomass and grain yield were evaluated at both per-row (g m⁻²) and field scales (kg ha⁻¹). Per-row values were calculated from three interior IWG rows × 3 m (9 linear m total), whereas field-scale values accounted for differences in IWG row density between monocultures (100% IWG rows) and intercrops (50% IWG rows).

Models with continuous or categorical stand age, with and without treatment × stand age interactions, were compared using maximum likelihood estimation, Akaike information criterion (AIC), and likelihood-ratio tests. Categorical stand age was retained based on model fit. Final models were fitted using restricted maximum likelihood (REML) with lme4, and fixed effects were assessed using Type III tests with Satterthwaite degrees of freedom in lmerTest (Kuznetsova et al., 2017). Estimated marginal means and pairwise contrasts were obtained using emmeans with Sidák adjustment. Model assumptions were evaluated using Shapiro-Wilk and Levene’s tests. IWG biomass at DDPSC FRS and alfalfa biomass at Planthaven were log(x + 1) transformed to address heteroscedasticity, with inference based on transformed models and back-transformed means reported for interpretation.

IWG lodging was analyzed separately for 2024 and 2025 because lodging was assessed once in 2024 at both sites, and at three sampling periods in 2025 at DDPSC FRS. In 2024, the effects of treatment on IWG lodging were evaluated within each site using linear models with treatment and block as fixed effects, whereas 2025 lodging analysis at DDPSC FRS included a linear mixed-effects model with treatment, sampling period, and their interaction as fixed effects and block as a random effect.

Total root length (TRL), root length density (RLD), specific root length (SRL), mean diameter, and root mass were analyzed using linear mixed-effects models with treatment, soil depth, and sampling position as fixed effects. Block and plot nested within block were included as random effects. Models were fitted using REML, and fixed effects were assessed using Type III tests with Satterthwaite degrees of freedom. Estimated marginal means and 95% confidence intervals were calculated for treatment × depth × position combinations, with Tukey-adjusted pairwise contrasts where relevant. Model assumptions were assessed using residual and Q-Q plots and the performance package (Lüdecke et al., 2021). Random-effect singularity and multicollinearity were evaluated using lme4 diagnostics and variance inflation factors, respectively. Statistical significance was assessed at α = 0.05.

## 3. RESULTS

### 3.1 Baseline soils

Baseline soil properties differed between sites (Fig. S1). Planthaven soil was strongly acidic (pH 4.68 ± 0.27), whereas DDPSC FRS soil was near neutral to slightly alkaline (pH 7.64 ± 0.08). DDPSC FRS soil also had greater CEC (19.9 ± 1.73 vs. 11.3 ± 0.61 cmolc kg⁻¹) and higher concentrations of K, Ca, Mg, total P, organic matter, and total organic carbon.

### 3.2 Weather conditions

Weather conditions were generally similar between sites within calendar years but varied across production years (Fig. S2, Table S1). Relative to the 1991-2020 normals, 2023 was drier at both sites, with annual precipitation deficits of 137 mm at Planthaven and 203 mm at DDPSC FRS, whereas 2024 was wetter than normal. Establishment was also drier than normal, particularly at Planthaven (−115 mm in September-November 2021), compared with DDPSC FRS (−45 mm in 2022; Table S1). During seedhead production (June-July), precipitation was above normal in all three production years at Planthaven but increased from 27 mm below normal in stand age 1 to 133 mm above normal in stand age 3 at DDPSC FRS. Temperatures during this period were generally near to above normal.

### 3.3 Per-row IWG aboveground biomass (g m⁻²)

At Planthaven, we observed that per-row aboveground IWG biomass was affected by treatment (F_2,24_ = 3.98, P = 0.032), stand age (F_2,24_= 33.32, P < 0.001), and their interaction (F_4,24_ = 3.15, P = 0.032; Table S2; Fig. 1A-C). IWG biomass increased from stand age 1 (285-352 g m⁻²) to stand age 2 (787-933 g m⁻²), with no significant treatment differences in either year. By stand age 3, IWG intercropped with alfalfa produced greater IWG biomass (1,019 g m⁻²) than the fertilized IWG monoculture plot (492 g m⁻²; P < 0.001), while IWG biomass in the unfertilized IWG monoculture was intermediate between the intercropped and fertilized IWG monoculture (761 g m⁻²).

**Figure 1.**
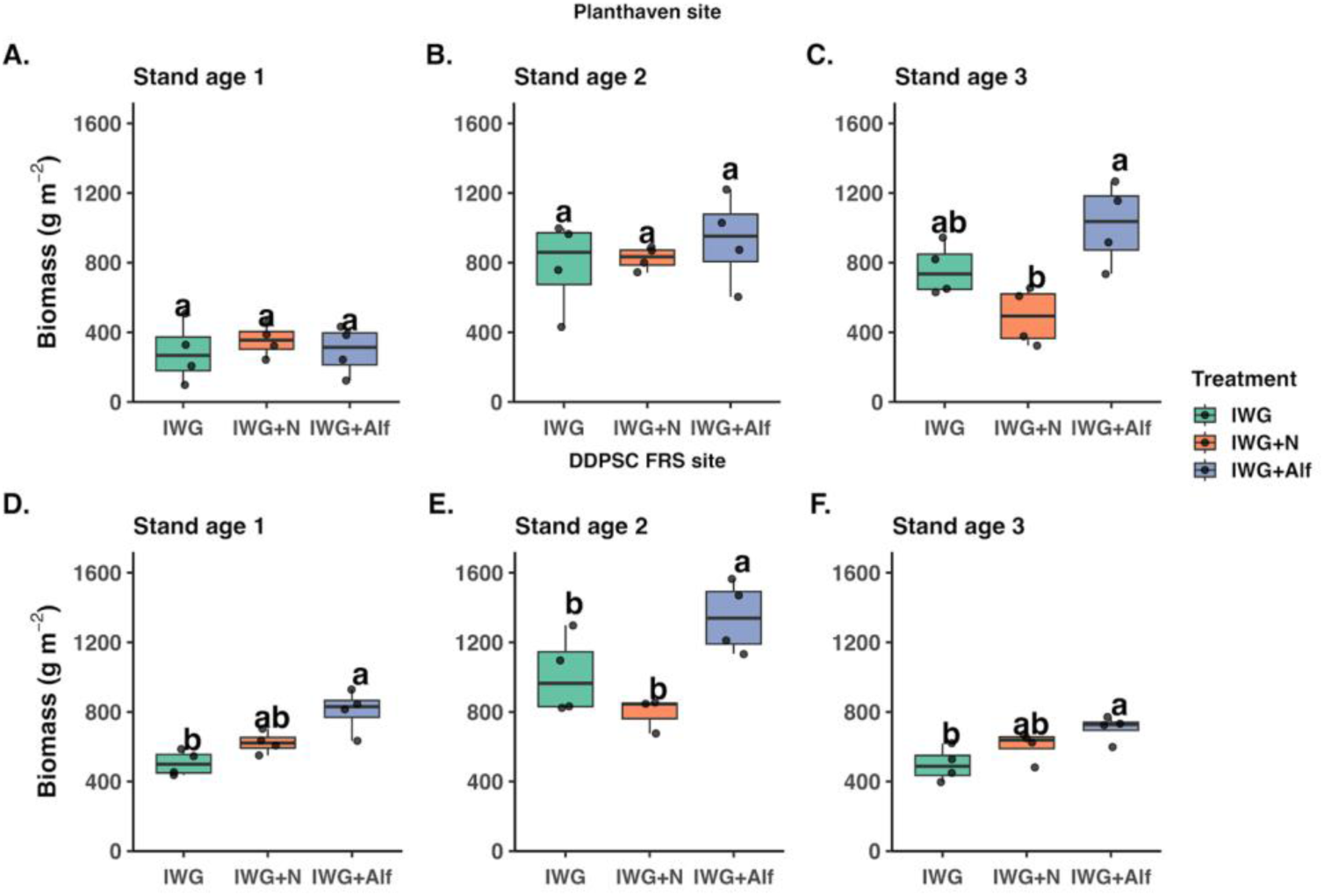
Intermediate wheatgrass (IWG) aboveground biomass at per-row level (g m⁻²) across three stand ages at the Planthaven site (A-C) and the DDPSC FRS site (D-F). Treatments include unfertilized IWG monoculture (IWG), nitrogen-fertilized IWG (IWG + N), and IWG intercropped with alfalfa (IWG + Alf). Different letters above treatments indicate significant differences within each stand age based on post-hoc comparisons (α = 0.05).

At DDPSC FRS, we observed that treatment (F_2,23.3_ = 20.17, P < 0.001) and stand age (F_2,23.3_ = 41.11, P < 0.001) affected per-row IWG aboveground biomass, whereas their interaction was not significant (F_4,23.2_ = 2.77, P = 0.051; Table S; Fig. 1D-F). IWG in the intercrop had the highest mean biomass across stand ages. In stand age 1, IWG biomass in the intercrop exceeded IWG biomass in the unfertilized IWG monoculture (798 vs. 502 g m⁻²; P < 0.001), but did not differ from the fertilized IWG (p = 0.089). Biomass peaked in stand age 2, with IWG biomass in the intercrop (1,332 g m⁻²) exceeding IWG biomass in both the unfertilized (994 g m⁻²; P = 0.037) and fertilized IWG monocultures (786 g m⁻²; P < 0.001). Although biomass declined in stand age 3, IWG biomass in the intercrop remained greater than IWG biomass in unfertilized IWG monoculture (703 vs. 491 g m⁻²; P = 0.009); fertilized IWG was intermediate and did not differ from either treatment.

### 3.4 Field-scale IWG aboveground biomass (kg ha⁻¹)

When we scaled IWG aboveground biomass to the field level at Planthaven, biomass was influenced by treatment (F_2,24_ = 10.20, P < 0.001), stand age (F_2,24_ = 33.66, P < 0.001), and their interaction (F_4,24_ = 2.93, P = 0.042; Table S3; Fig. 2A-C). No significant treatment differences were detected in stand age 1. By stand age 2, unfertilized and fertilized IWG monocultures produced greater field-scale IWG biomass (7,871 and 8,249 kg ha⁻¹, respectively) than the intercrop (4,663 kg ha⁻¹; P ≤ 0.007). In stand age 3, the unfertilized IWG monoculture maintained greater IWG biomass (7,610 kg ha⁻¹) than fertilized IWG monoculture (4,917 kg ha⁻¹; P = 0.027) and the intercrop (5,096 kg ha⁻¹; P = 0.042), which did not differ from one another.

**Figure 2.**
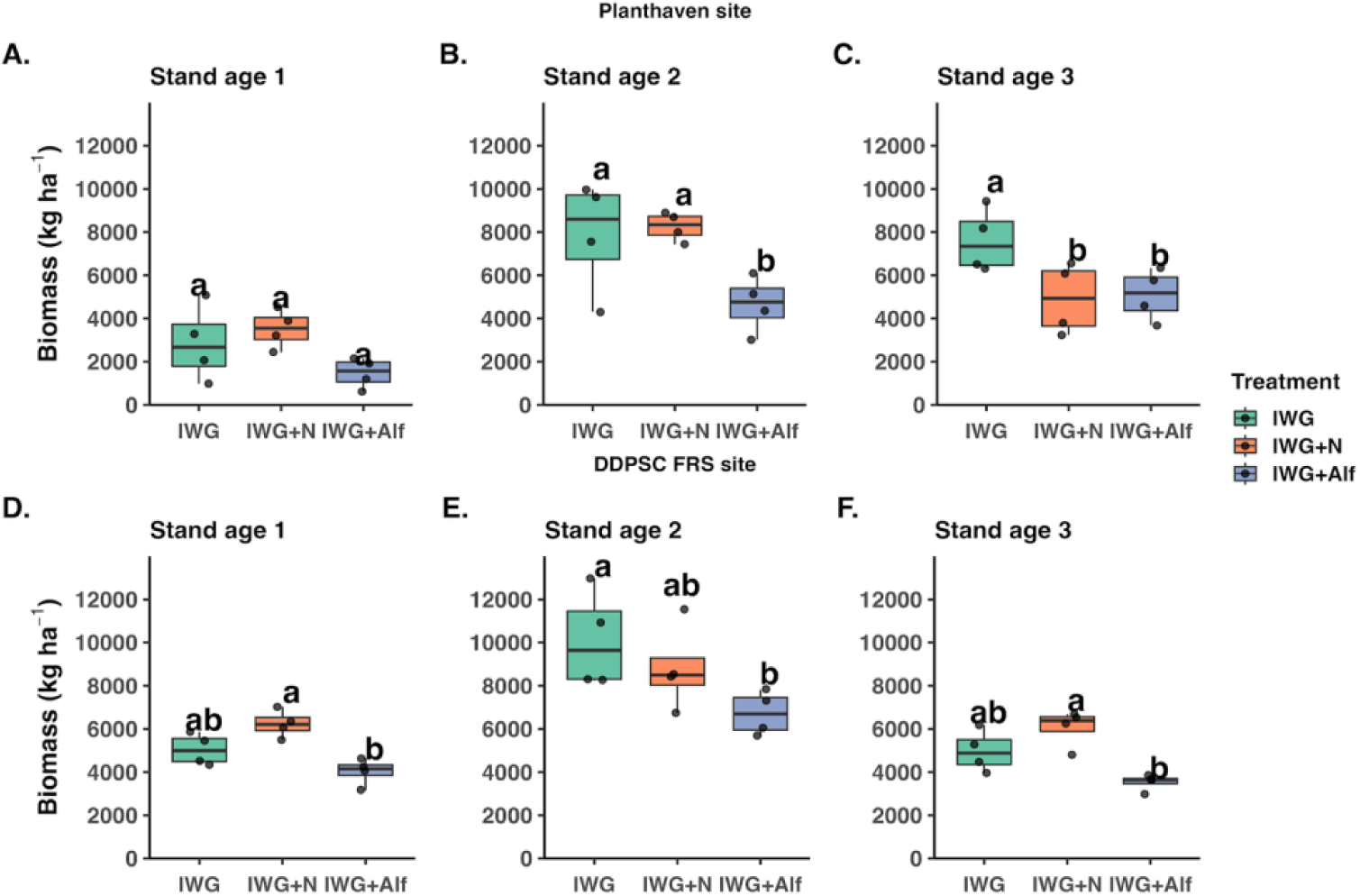
Intermediate wheatgrass (IWG) aboveground biomass at field level (kg ha⁻¹) across three stand ages at the Planthaven site (A-C) and the DDPSC FRS site (D-F). Treatments include unfertilized IWG, nitrogen-fertilized IWG (IWG + N), and IWG intercropped with alfalfa (IWG + Alf). Different letters above treatments indicate significant differences within each stand age based on post-hoc comparisons (α = 0.05).

At DDPSC FRS, treatment (F_2,24_ = 12.72, P < 0.001) and stand age (F_2,24_ = 35.57, P < 0.001) affected field-scale IWG biomass, but their interaction was not significant (F_4,24_ = 1.69, P = 0.185; Table S3; Fig. 2D-F). Averaged across stand ages, unfertilized and fertilized IWG monocultures produced greater IWG biomass (6,718 and 7,043 kg ha⁻¹, respectively) than the intercrop (4,759 kg ha⁻¹; P ≤ 0.002), with no difference between the monocultures. Across treatments, IWG biomass peaked in stand age 2 (8,555 kg ha⁻¹), exceeding stand ages 1 (5,107 kg ha⁻¹) and 3 (4,857 kg ha⁻¹; both P < 0.001).

### 3.5 Per-row IWG grain yield (g m⁻²)

We observed that at Planthaven, per-row IWG grain yield was affected by treatment (F_2,27_ = 4.72, P = 0.017), stand age (F_2,27_ = 7.48, P = 0.003), and their interaction (F_4,27_= 4.20, P = 0.009; Table S2; Fig. 3A-C). Grain yield increased from stand age 1 (40.1-67.3 g m⁻²) to stand age 2 (79.9-118.9 g m⁻²), with no significant treatment differences in either year. By stand age 3, IWG intercropped with alfalfa produced greater IWG grain yield (133.0 g m⁻²) than the fertilized IWG monoculture (36.6 g m⁻²; P < 0.001), while IWG biomass in the unfertilized monoculture was intermediate (83.2 g m⁻²).

**Figure 3.**
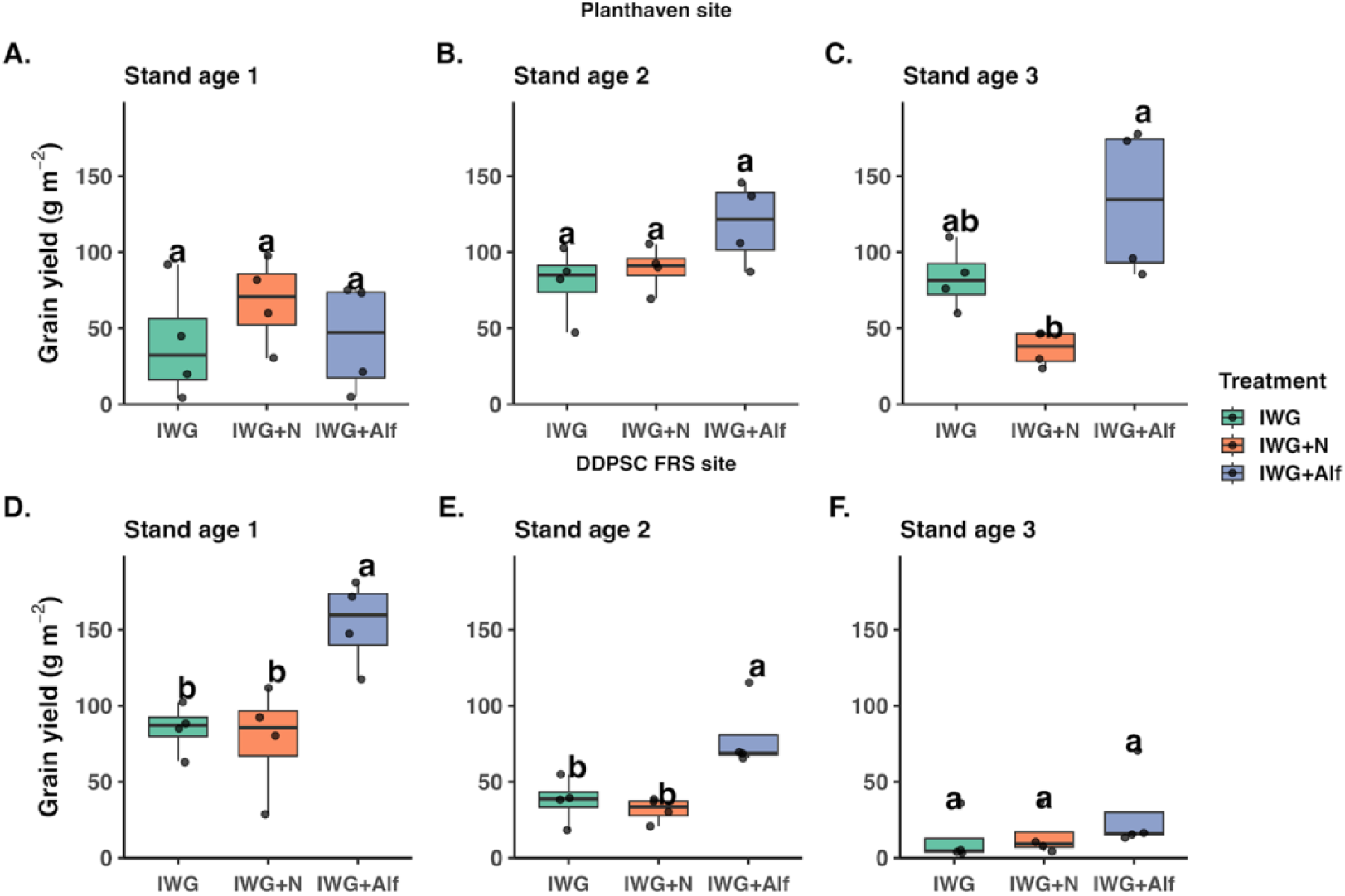
IWG grain yield at per-row level (g m⁻²) across three stand ages at Planthaven (A-C) and DDPSC FRS (D-F). Treatments: IWG, IWG + N, and IWG + Alf. Boxplots show medians and variability; letters denote significant differences within each stand age (α = 0.05).

At DDPSC FRS, treatment (F_2,24_ = 19.20, P < 0.001) and stand age (F_2,24_ = 56.90, P < 0.001) affected per-row grain yield, whereas their interaction was not significant (F_4,24_ = 2.71, P = 0.054; Table S2; Fig. 3D-F). Averaged across stand ages, IWG intercropped with alfalfa produced greater grain yield (87.6 g m⁻²) than unfertilized (45.0 g m⁻²) and fertilized IWG monocultures (41.5 g m⁻²; both P < 0.001), with no difference between the monoculture treatments. Across all treatments, grain yield declined from 105.8 g m⁻² in stand age 1 to 49.7 g m⁻² in stand age 2 and 18.6 g m⁻² in stand age 3, with all stand ages differing significantly over years (P ≤ 0.003).

### 3.6 Field-scale IWG grain yield (kg ha⁻¹)

Field-scale IWG grain yield followed contrasting trajectories at the two sites (Fig. 4). At Planthaven, yield varied with stand age (F_2,24_ = 6.30, P = 0.006), and the significant treatment × stand-age interaction (F_4,24_ = 3.88, P = 0.014) reflected changes in treatment responses over time. The overall treatment effect was not significant (F_2,24_ = 2.20, P = 0.133; Table S3). Fertilized IWG monocultures initially produced greater IWG grain yield than the intercrop (673 vs. 219 kg ha⁻¹; P = 0.029), with unfertilized IWG monocultures intermediate (401 kg ha⁻¹). By stand age 2, grain yields had increased across all treatments (595-893 kg ha⁻¹), and no significant differences among treatments were detected. This pattern shifted in stand age 3, when unfertilized IWG monoculture produced greater yield (832 kg ha⁻¹) than fertilized IWG monoculture (366 kg ha⁻¹; P = 0.025), while the intercrop remained intermediate (665 kg ha⁻¹).

**Figure 4.**
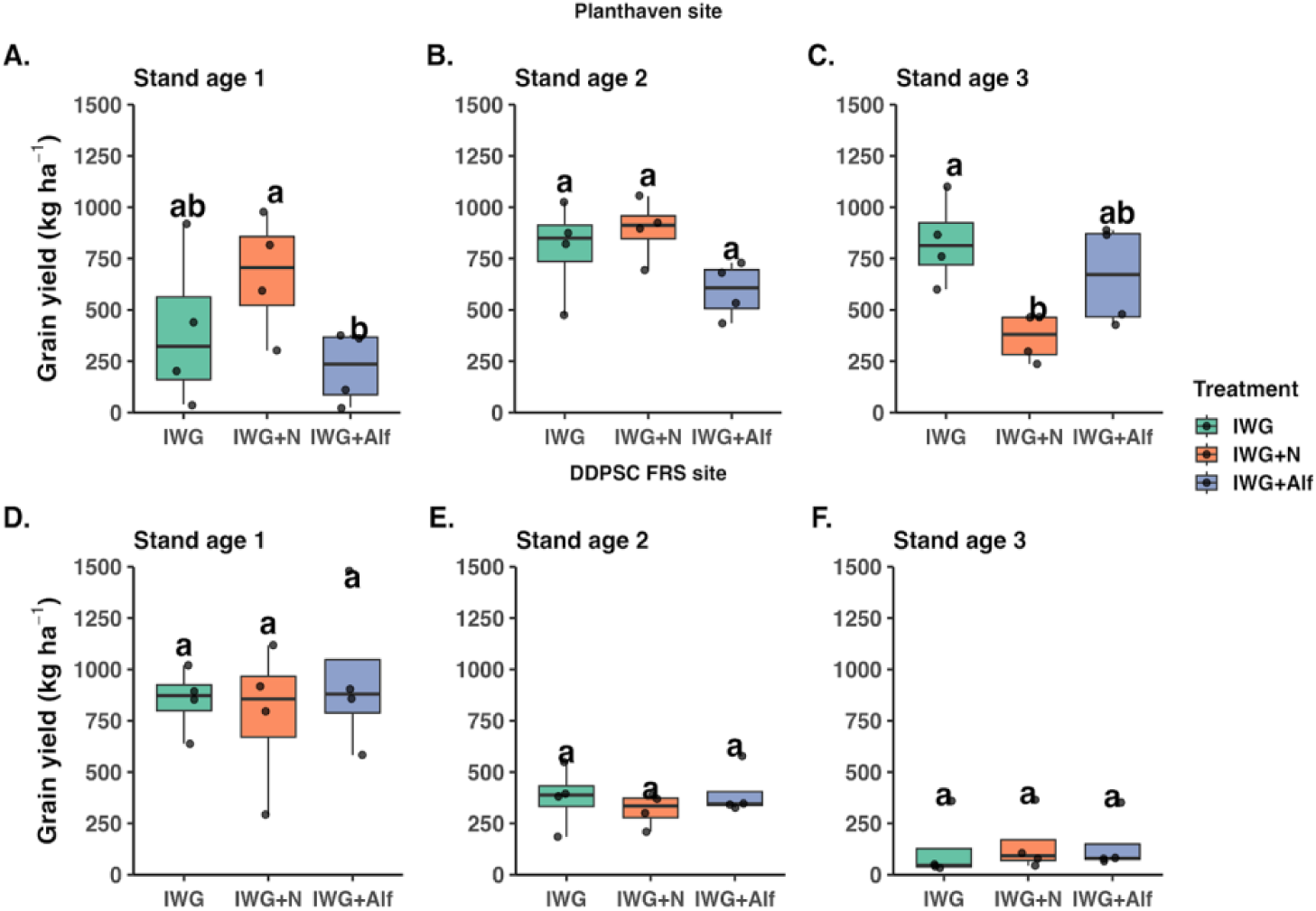
Intermediate wheatgrass (IWG) grain yield at field level (kg ha⁻¹) across three stand ages at the Planthaven site (A-C) and DDPSC FRS site (D-F). Treatments include unfertilized IWG, nitrogen-fertilized IWG (IWG + N), and IWG intercropped with alfalfa (IWG + Alf). Different letters above treatments indicate significant differences within each stand age based on post-hoc comparisons (α = 0.05).

At DDPSC FRS, grain yield declined markedly with stand age, irrespective of treatment (F_2,24_ = 46.74, P < 0.001; Table S3; Fig. 4D-F). Yields fell from 781-956 kg ha⁻¹ in stand age 1 to 316-398 kg ha⁻¹ in stand age 2, reaching only 121-149 kg ha⁻¹ by stand age 3. Neither the treatment main effect (F_2,24_= 0.61, P = 0.551) nor its interaction with stand age (F_4,24_ = 0.25, P = 0.908) was significant. Notably, the intercrop maintained field-scale IWG grain yield comparable to the fertilized and unfertilized IWG monocultures despite having only half as many IWG rows.

### 3.7 Alfalfa aboveground biomass

At Planthaven, per-row alfalfa aboveground biomass (g m⁻²) varied with treatment (F_1,18_ = 14.19, P = 0.001) and stand age (F_2,18_ = 34.78, P < 0.001), but there was no significant interaction between the two (F_2,18_ = 0.51, P = 0.609; Table S2). Alfalfa monoculture produced greater aboveground biomass than intercropped alfalfa, although the difference was significant only in stand age 3 (394 vs. 39 g m⁻²; P = 0.025; Fig. S3). At DDPSC FRS, treatment (F_1,15_ = 46.55, P < 0.001) and stand age (F_2,15_ = 87.10, P < 0.001) also affected per-row alfalfa aboveground biomass, without a significant interaction (F_2,15_ = 2.78, P = 0.094; Table S2). Monoculture biomass exceeded intercrop biomass in stand ages 2 and 3 (P < 0.004; Fig. S3).

At the field scale (kg ha⁻¹), alfalfa monocultures produced greater biomass than intercropped alfalfa at both sites (Fig. S4). At Planthaven, field-scale alfalfa aboveground biomass was affected by treatment (F_1,15_ = 98.38, P < 0.001), stand age (F_2,15_ = 43.71, P < 0.001), and their interaction (F_2,15_ = 15.84, P < 0.001; Table S4). Aboveground biomass increased in both treatments from stand age 1 to 2 before declining in stand age 3, when monoculture alfalfa produced 3,885 kg ha⁻¹ compared with 154 kg ha⁻¹ in the intercrop (P < 0.001). At DDPSC FRS, field-scale alfalfa aboveground biomass was also affected by treatment (F_1,15_ = 195.73, P < 0.001) and stand age (F_2,15_ = 64.62, P < 0.001), but not their interaction (F_2,15_ = 3.49, P = 0.057; Table S4). Biomass declined with stand age in both treatments, and monoculture alfalfa consistently exceeded intercropped alfalfa (P < 0.001 across stand ages; Fig. S4).

### 3.8 IWG Lodging

Lodging was generally greatest in fertilized IWG monocultures, although treatment responses varied by site and stand age (Table S4; Fig. S5). At Planthaven in stand age 3, the estimated proportion of lodged IWG was 82.5% in fertilized monocultures, compared with 43.8% in the intercrop and 8.7% in unfertilized monocultures (F_2,6_ = 16.64, P = 0.004). We observed a similar pattern at DDPSC FRS in stand age 2, where lodging was 60.0% in fertilized monoculture compared with 4.3% in the intercrop and 3.3% in unfertilized IWG monoculture (F_2,6_ = 71.55, P < 0.001). At DDPSC FRS in stand age 3, no lodging was observed in May, but lodging increased later in the season in fertilized and intercropped IWG while remaining low in unfertilized IWG (treatment × observation time: F_4,24_ = 16.27, P < 0.001).

### 3.9 Root responses to treatment, soil depth, and sampling position

Root traits varied primarily with soil depth, with comparatively limited responses to treatments at either site (Table S5; Figs. 5-6). TRL, RLD, and root mass were generally greatest in the surface soil (0-20 cm). Depth significantly affected TRL at Planthaven (F_2,117_ = 6.63, P = 0.002) and DDPSC FRS (F_2,117_ = 50.94, P < 0.001), as well as RLD at both sites (Planthaven: F_2,117_ = 38.75, P < 0.001; DDPSC FRS: F_2,112.1_ = 128.56, P < 0.001). Root mass also declined with depth at both sites.

**Figure 5.**
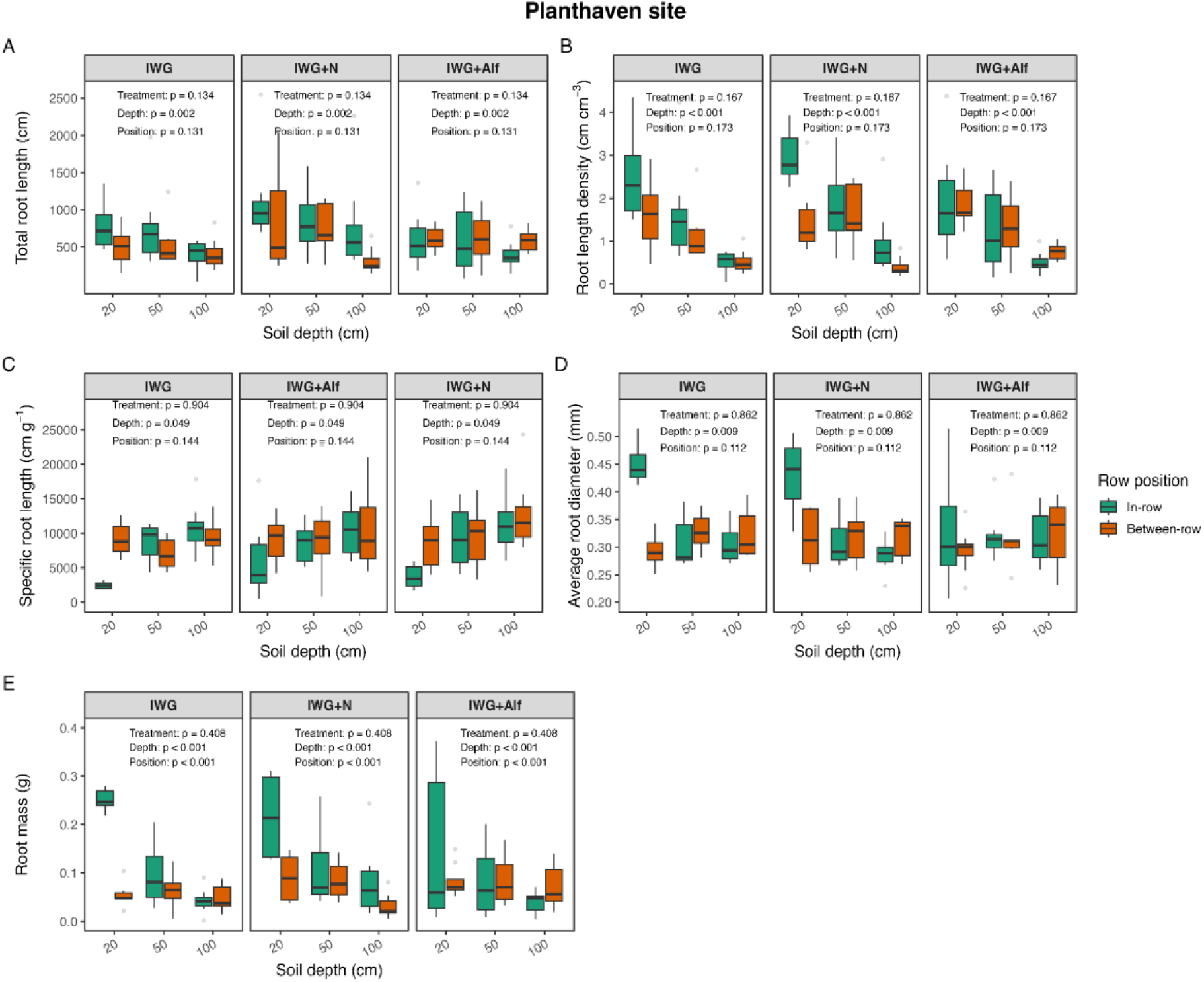
Root responses of IWG to treatment, soil depth, and row position at the Planthaven site. Panels show (A) total root length (TRL), (B) root length density (RLD), (C) specific root length (SRL), (D) average root diameter, and (E) root mass measured at three soil depths (0-20, 20-50, and 50-100 cm) for unfertilized IWG (IWG), nitrogen-fertilized IWG (IWG+N), and IWG intercropped with alfalfa (IWG+Alf). Colors denote sampling position (green = in-row; orange = between-row). Reported *p*-values summarize Type III mixed-effects ANOVA results for main effects of treatment, depth, and position.

**Figure 6.**
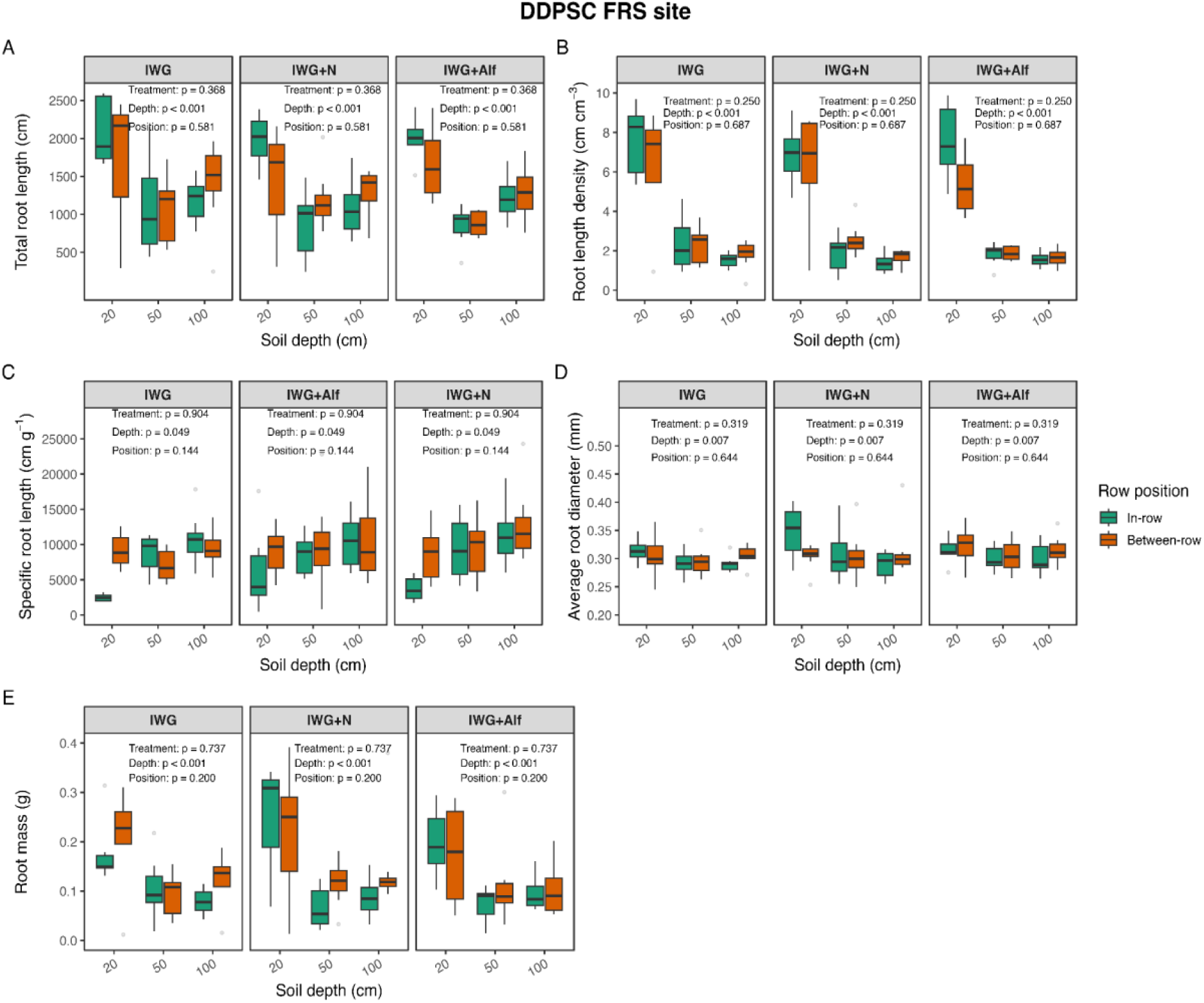
Root trait responses of IWG to treatments, soil depth, and row position at the DDPSC FRS site. Panels show (A) total root length (TRL), (B) root length density (RLD), (C) specific root length (SRL), (D) average root diameter, and (E) root mass measured at three soil depths (0-20, 20-50, and 50-100 cm) for unfertilized IWG (IWG), nitrogen-fertilized IWG (IWG+N), and IWG intercropped with alfalfa (IWG+Alf). Colors indicate sampling position (green = in-row; orange = between-row). Reported *p*-values summarize Type III mixed-effects ANOVA results for treatment, depth, and position effects.

Although treatment had little influence on overall root traits, some differences emerged in their spatial distribution. At Planthaven, TRL showed a treatment × sampling-position interaction (F_2,117_ = 3.21, P = 0.044). Fertilized IWG tended to have greater TRL within rows, whereas the intercrop showed the opposite pattern. Root diameter also showed a treatment × depth interaction at Planthaven (F_4,117_ = 3.36, P = 0.012), with fertilized IWG having thicker roots than the intercrop in the surface layer (P = 0.045). No treatment differences in root diameter were detected at greater depths.

At Planthaven, root mass showed a depth × position interaction (F_2,124_ = 20.40, P < 0.001), with greater in-row than between-row root mass in the surface layer (P < 0.001). Root diameter also varied with depth and position at both sites (Planthaven: F_2,117_ = 13.95, P < 0.001; DDPSC FRS: F_2,115.9_ = 3.89, P = 0.023). In-row root diameter generally declined with depth, whereas between-row diameter remained more uniform.

## 4. DISCUSSION

### 4.1 IWG biomass and grain yield responses to fertilization and intercropping varied with stand age and measurement scales

Alfalfa intercropping maintained IWG vegetative biomass and grain yield at levels comparable to, and in some cases greater than, nitrogen-fertilized monocultures. Although nitrogen fertilization increased vegetative biomass in some site-years, particularly at DDPSC FRS (Fig. 1D-F), these increases did not translate into consistent grain yield (Fig. 3D-F). Previous studies have similarly reported variable IWG grain yield responses to nitrogen fertilization and legume intercropping (Jungers et al. 2017; Fernandez et al. 2020; Pinto et al. 2024; McKenna et al. 2026). Our findings extend these observations by demonstrating that treatment responses varied with stand age even between nearby sites.

The relative performance of intercropped and monoculture IWG also depended on measurement scale. On a per-row basis, IWG in the intercrop produced vegetative biomass and grain yield comparable to, and in some site-years greater than, IWG in the fertilized monoculture. At the field scale, intercrop grain yield was comparable to that of fertilized IWG in several site-years despite having half as many IWG rows. Greater yield per IWG row could reflect reduced competition, interactions with alfalfa, or both. Similar benefits of cereal-legume intercropping have been associated with changes in competition, complementary resource use, and biological nitrogen fixation (Jensen et al., 2020; Crews et al., 2022; Raza et al., 2023; Luo et al., 2024). However, because our study did not include a density-matched IWG monoculture or directly measure nitrogen transfer, we cannot determine which mechanisms contributed to the observed differences.

Treatment responses also shifted as stands matured. At Planthaven, per-row vegetative biomass and grain yield of fertilized IWG declined between stand ages 2 and 3, whereas per-row biomass of intercropped IWG continued to increase, and grain yield remained relatively high. At DDPSC FRS, vegetative biomass increased from stand age 1 to 2 before declining in stand age 3, while grain yield declined substantially after stand age 1 across treatments (Figs. S6A, S6B). These contrasting patterns are consistent with previous studies showing that IWG aboveground biomass and grain yield change with stand age (Zhen et al., 2024; Ding et al., 2025). Changes in plant competition, nutrient cycling, and resource availability may contribute to these responses as perennial stands mature (Jungers et al., 2015; Sharma et al., 2022; Louarn et al., 2024). The increasing advantage of alfalfa-intercropped IWG over fertilized IWG at Planthaven suggests that treatment responses may shift as stands mature, although the influence of alfalfa cannot be separated from reduced IWG density.

Lodging provides additional context for the lack of a consistent grain yield response to nitrogen fertilization. At Planthaven in stand age 3, lodging was greatest in fertilized IWG (82.5%), followed by intercropped IWG (43.8%) and unfertilized IWG (8.7%). Fertilized IWG also had greater lodging than the other treatments at DDPSC FRS in stand ages 2 and 3 (Fig. S5; Table S4). This pattern is consistent with studies showing that increased nitrogen availability can increase lodging susceptibility in cereal crops (Wu et al., 2019; Lu et al., 2025). However, lodging cannot be identified as the cause of the observed grain yield patterns because lodged stems were included in our harvests and yield components were not quantified. Together, our findings suggest that intercropping can maintain IWG productivity comparable to nitrogen fertilization in some environments and stand ages, but further studies incorporating density-matched monocultures and direct measurements of nitrogen dynamics are needed to identify the mechanisms responsible.

### 4.2 Root traits were structured primarily by soil depth rather than management

In contrast to the aboveground responses of vegetative biomass and grain yield to nitrogen fertilization and alfalfa intercropping (here and in McKenna et al. 2026), root traits were influenced primarily by soil depth rather than treatment, consistent with previous studies (e.g., Sprunger et al. 2017; Clément et al. 2022; Sakiroglu et al. 2020; Schenk and Jackson 2002). RLD and root mass were concentrated primarily in the 0-20 cm soil layer, while TRL and average root diameter also varied with depth. Although treatment main effects were generally weak, some spatial responses were evident. At Planthaven, the significant treatment × sampling-position interaction for TRL reflected a tendency for fertilized IWG to have greater root length within planting rows, whereas the alfalfa intercrop showed the opposite numerical pattern. However, individual in-row versus between-row contrasts were not significant after adjustment, so these patterns should be interpreted cautiously. Previous studies have reported that root proliferation can respond to localized resource availability and neighbouring plants (Hodge, 2004; Hauggaard-Nielsen and Jensen, 2005; Zhang et al., 2020), but our measurements did not distinguish whether the observed spatial patterns resulted from nutrient distribution, interspecific interactions, or other factors.

The limited treatment effects on root traits may also reflect the spatial and temporal constraints of the soil-core sampling method used here. Root traits were measured at a single sampling period during the second stand age, providing a spatial snapshot rather than a measure of seasonal root production, turnover, or activity. Soil cores also sample only a small portion of the root system (Taylor et al., 2013), so treatment effects occurring at other times or through more dynamic belowground processes may have gone undetected (Freschet et al., 2021). Repeated measurements or complementary approaches such as minirhizotron imaging could reveal management responses not captured by a single soil-core sampling.

### 4.3 Site context shapes IWG biomass and grain responses at a local scale

Previous studies have reported site- and year-dependent responses to management in IWG systems (Jungers et al., 2017; Tautges et al., 2018; Reilly et al., 2022; Cassani et al., 2025; Mckenna et al. 2026). Our study extends these findings by showing that contrasting management responses can also occur between nearby sites within the same climatic region.

Site-specific differences may have influenced treatment response. Both Planthaven and the DDPSC FRS were farmed conventionally with corn and soybeans for decades prior to their use in this study, but unfortunately, we do not have yield data from those years for those crops. Interestingly, baseline soil properties of our two study sites reveal important differences that may have impacted IWG growth and production (Fig. S1). Planthaven had strongly acidic, poorly drained, high-clay soils, whereas DDPSC FRS had better-drained soils with higher pH and nutrient availability. These contrasting soil environments may have contributed to differences in crop establishment and subsequent responses to fertilization and intercropping. Consistent with the importance of site context, Cassani et al. (2025) also found that edaphic and climatic conditions explained more variation in IWG grain yield than N fertilization across six Midwestern sites. These findings underscore the relevance of local soil properties for IWG site selection and productivity expectations.

Establishment conditions and interannual weather may also have contributed to differences between sites. Because the sites were established in different years, site and establishment year cannot be separated in our study. Both sites received below-normal precipitation during fall establishment, but the deficit was greater at Planthaven in 2021 (Fig. S2; Table S1). Precipitation also varied during production years, and its timing may be particularly important for IWG yield (Demers et al., 2026). Precipitation effects in our study may have depended on both timing and magnitude and may have interacted with local soil conditions, particularly at Planthaven, where high clay content and poor drainage could have amplified the effects of wet periods.

## 5. CONCLUSION

Alfalfa intercropping maintained or enhanced IWG grain yield in several site-years relative to nitrogen fertilization, although IWG responses to nitrogen and alfalfa intercropping depended strongly on local site conditions and stand age. Future research should evaluate a broader diversity of legume species and mixtures, optimize legume-IWG density and spatial arrangements, and directly quantify nitrogen transfer and resource use. Such approaches will help identify intercrop designs that sustain IWG yield while reducing reliance on synthetic N fertilizer.

## Supporting information

Supplementary site information, statistical analyses of IWG biomass and grain yield, lodging, and root traits.

## AUTHOR CONTRIBUTIONS

This study was conceptualized by TM, TC, and AM. Methodology was developed by TM, MH, TC, and AM. Investigation was conducted by KO, MH, SM, NW, TT, and AM. Data were curated by KO, TM, and SM, while formal analysis was performed by KO and TM. Project administration was carried out by TM, MH, and AM. Resources were provided by TM and AM, and funding was acquired by TC and AM. Supervision was provided by KO, TM, MH, TC, and AM. Data visualization was completed by KO, while validation was performed by TM and AM. KO wrote the original draft. All authors contributed to reviewing and editing the manuscript and approved the final version.

## ACKNOWLEGMENTS

We thank Sam Selby, Jack Braley, Isabella Vegara, Danielle Hopkins, Deva Raj Khanal, Biva Chapagain, Karen Schmiedeler, Stella Woeljen, Allie Schenewerk, Paige Bostic, Gillian Doty, Emma Vagelos, Azaria Covington, Maxwell Look, Zoraya Piedra, Matt McNeese, Mathew Rubin, Madeline DuBois, and Laura Van der Pol for their assistance with field establishment and maintenance, field sampling and harvesting, sample processing, and laboratory analyses. We also thank the staff and research personnel at Planthaven Farms and the Donald Danforth Plant Science Center Field Research Site for their support of the field experiments.

## Abbreviations

CEC: cation exchange capacity
IWG: intermediate wheatgrass
IWG+Alf: intermediate wheatgrass intercropped with alfalfa
IWG+N: nitrogen-fertilized intermediate wheatgrass
OM: organic matter
RLD: root length density
SRL: specific root length
TOC: total organic carbon
TRL: total root length.

