## Supplementary site information, statistical analyses of IWG biomass and grain yield, lodging, and root traits. for "Alfalfa intercropping maintains intermediate wheatgrass grain yield relative to nitrogen fertilization across site conditions and stand ages"

#### Supplementary Figures:

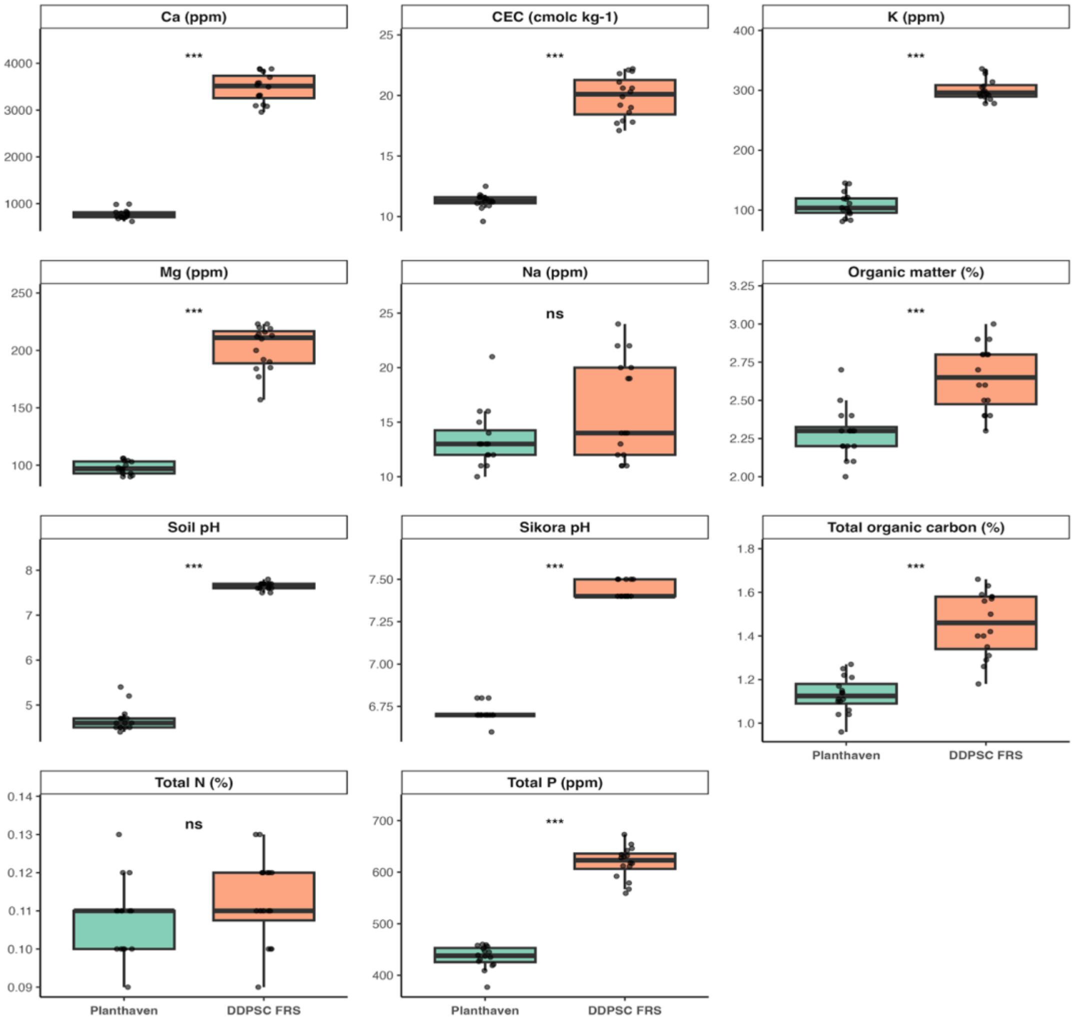


**Figure S1.** Baseline soil properties at the Planthaven and DDPSC FRS sites prior to treatment implementation. Boxplots show the distribution of soil calcium (Ca), cation exchange capacity (CEC), potassium (K), magnesium (Mg), sodium (Na), organic matter, soil pH, Sikora pH, total organic carbon (TOC), total nitrogen (N), and total phosphorus (P) across plots at each site (n = 16 plots per site). *** indicates BH-adjusted $P < 0.001$ and ns indicates no significant difference ($P \geq 0.05$).

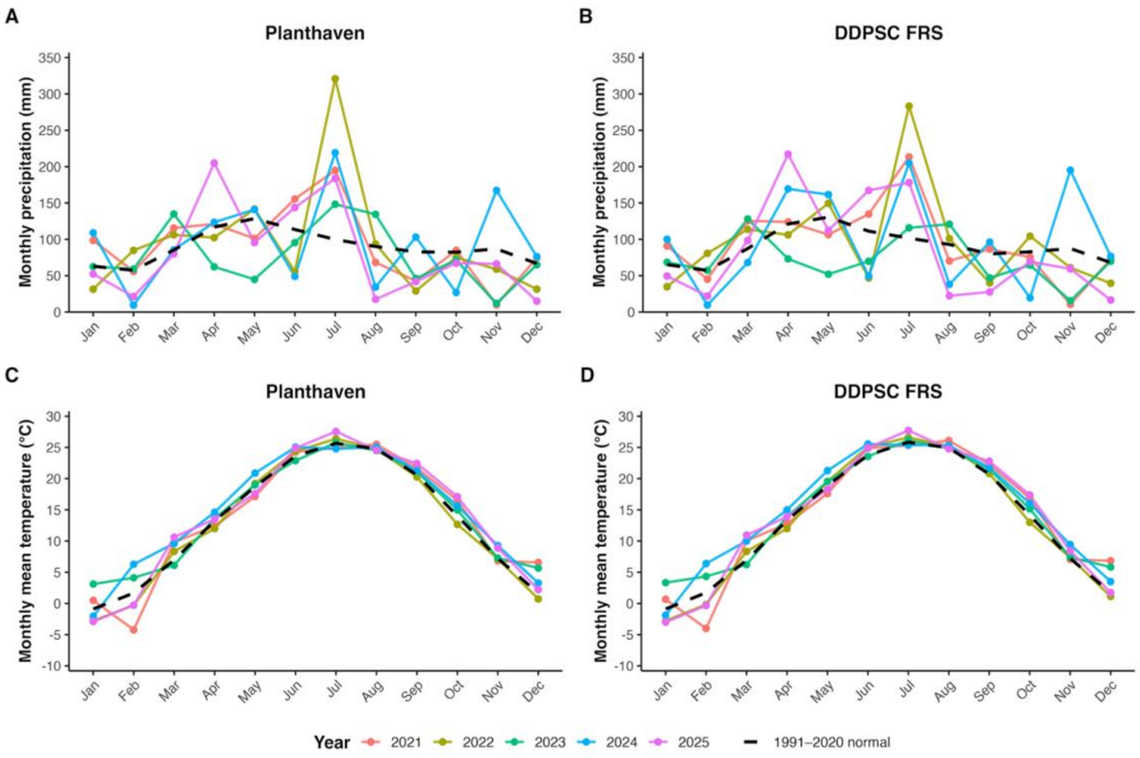


**Figure S2.** Monthly precipitation and mean air temperature at Planthaven and DDPSC FRS from 2021 to 2025 relative to 1991-2020 climate normals. Monthly precipitation at (A) Planthaven and (B) DDPSC FRS, and monthly mean air temperature at (C) Planthaven and (D) DDPSC FRS. Colored lines represent individual years (2021-2025), and dashed black lines represent the corresponding 1991-2020 monthly climate normals.

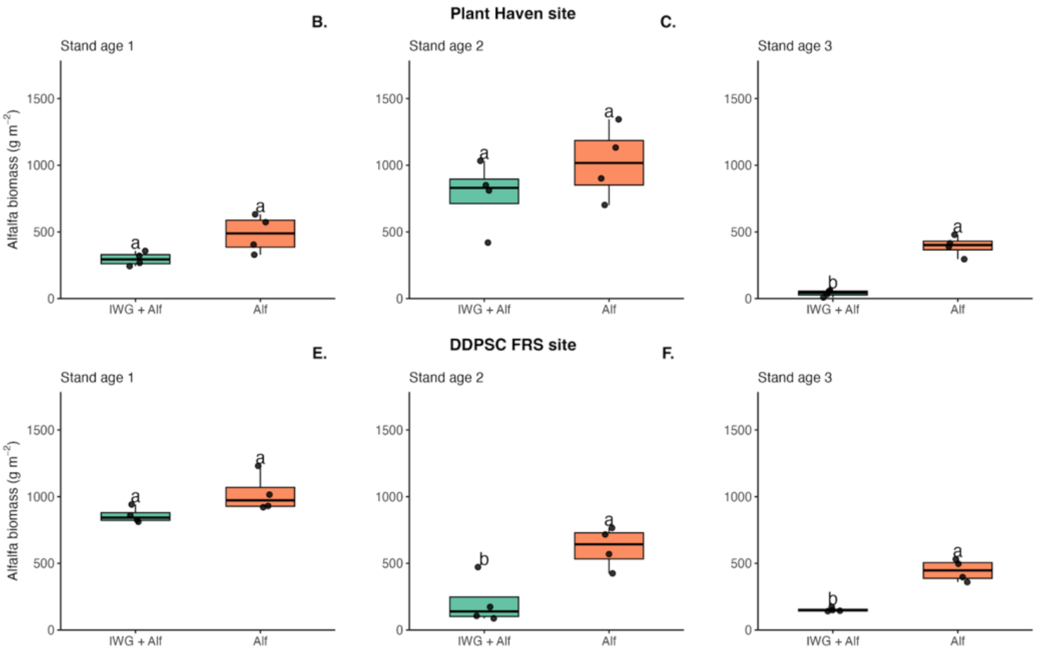


**Figure S3.** Alfalfa biomass (g m⁻2) across three stand ages at the Plant Haven site (A-C) and the DDPSC FRS site (D-F). Treatments include alfalfa monoculture and alfalfa grown in biculture with intermediate wheatgrass (IWG). Different letters above treatments indicate significant differences within each stand age based on post-hoc comparisons (α = 0.05).

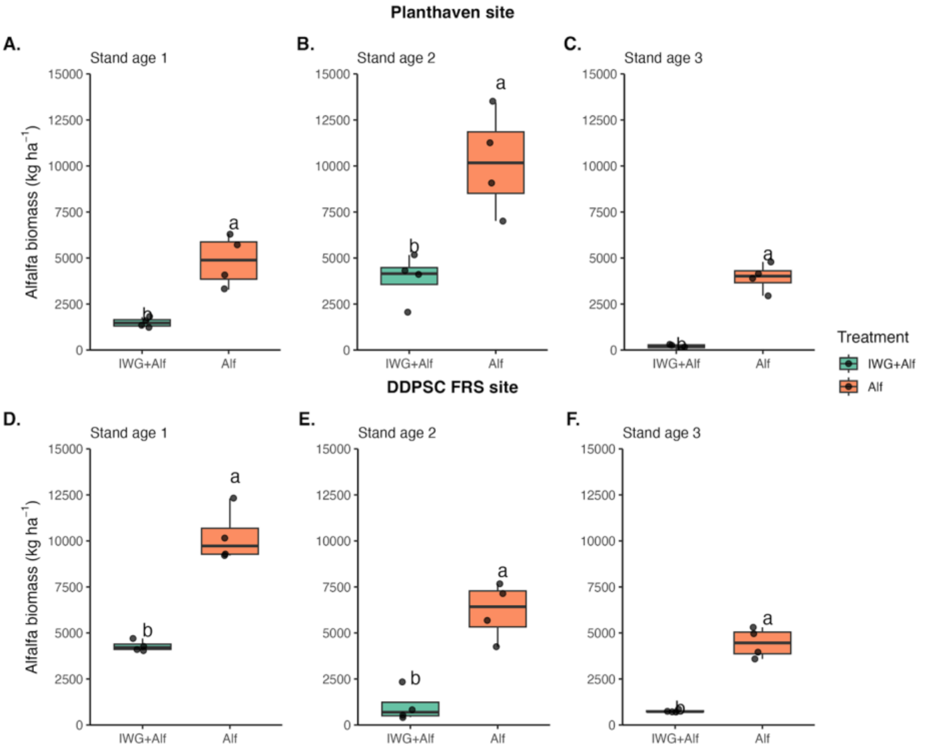


**Figure S4.** Alfalfa biomass (kg ha$^{-1}$) across three stand ages at the Plant Haven site (A-C) and the DDPSC FRS site (D-F). Treatments include alfalfa monoculture and alfalfa grown in biculture with intermediate wheatgrass (IWG). Different letters above treatments indicate significant differences within each stand age based on post-hoc comparisons (α = 0.05).

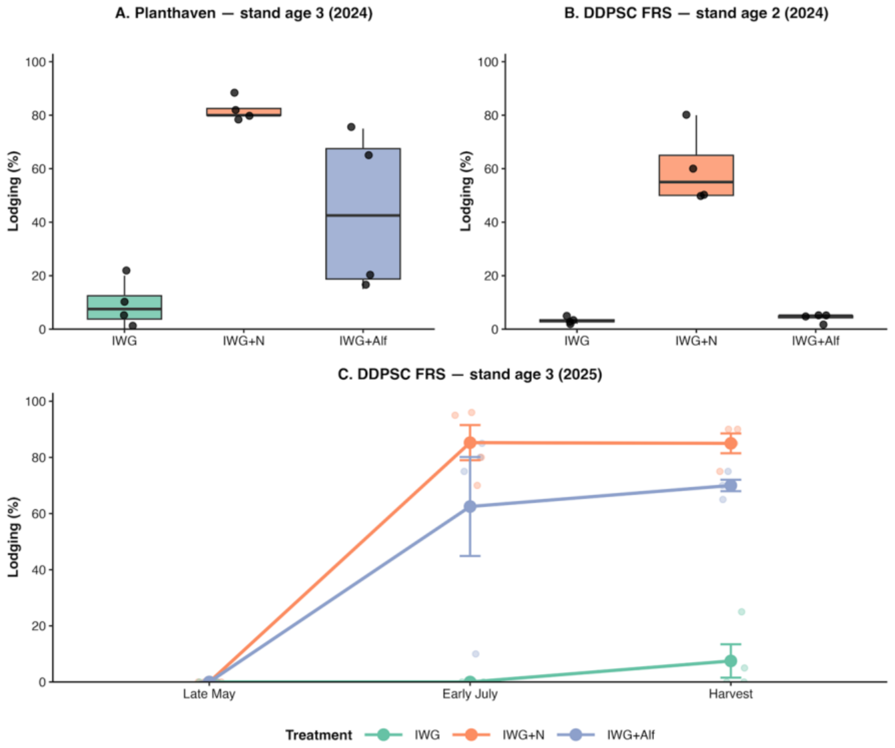


**Figure S5**. Lodging of IWG under different treatments at Planthaven and DDPSC FRS. (Panel A) Lodging at Planthaven in stand age 3 (2024). (B) Lodging at DDPSC FRS in stand age 2 (2024). (C) Seasonal progression of lodging at DDPSC FRS in stand age 3 (2025), assessed in May, July, and immediately before harvest. Treatments were unfertilized IWG monoculture (IWG), nitrogen-fertilized IWG monoculture (IWG+N), and IWG intercropped with alfalfa (IWG+Alf)

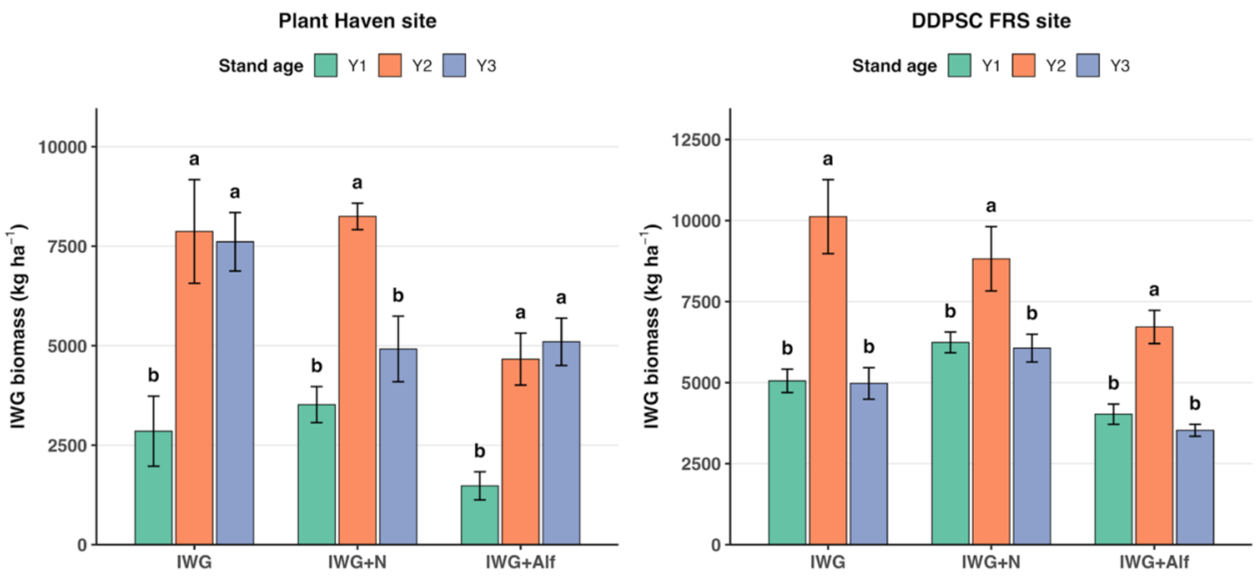


**Figure S6A.** Intermediate wheatgrass biomass (kg ha$^{-1}$) across three stand ages (Y1-Y3) within each cropping treatment at the Planthaven (left) and DDPSC FRS sites (right). Treatments included unfertilized IWG monoculture (IWG), nitrogen-fertilized IWG monoculture (IWG+N), and IWG intercropped with alfalfa (IWG+Alf). Bars represent treatment means ± SE. Different letters indicate significant differences among stand ages within each treatment and site based on Tukey-adjusted pairwise comparisons (α = 0.05).

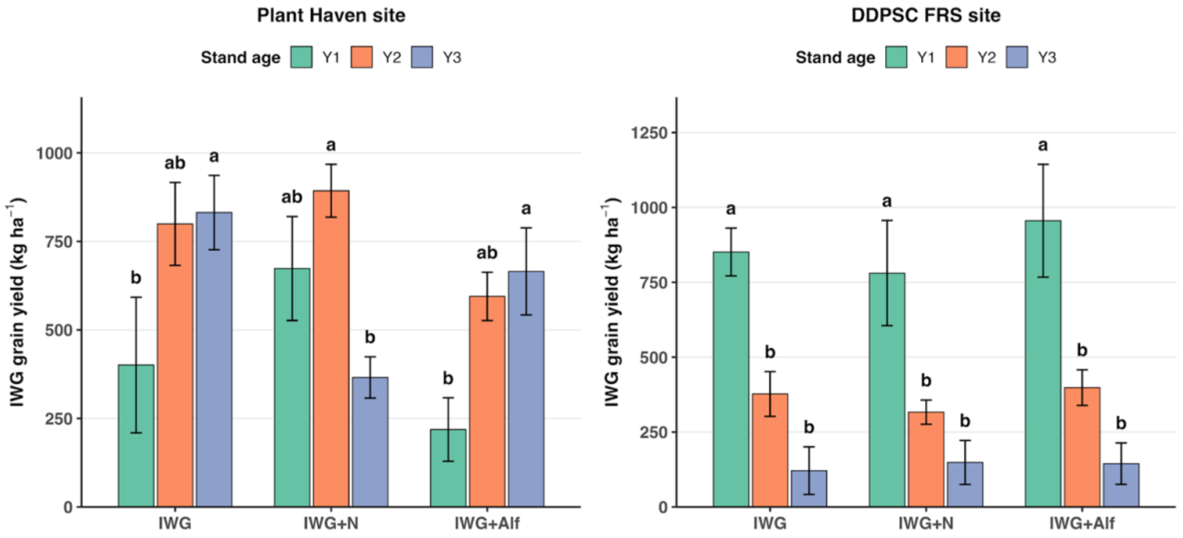


**Figure S6B.** Intermediate wheatgrass grain yield (kg ha$^{-1}$) across three stand ages (Y1-Y3) within each cropping treatments at the Planthaven (left) and DDPSC FRS sites (right). Treatments included unfertilized IWG monoculture (IWG), nitrogen-fertilized IWG monoculture (IWG+N), and IWG intercropped with alfalfa (IWG+Alf). Different letters indicate significant differences among stand ages within each treatment and site based on Tukey-adjusted pairwise comparisons (α = 0.05).

### Supplementary Tables:

#### Table S1. Precipitation and temperature during establishment and reproductive development at Planthaven and DDPSC FRS relative to 1991-2020 climate normals.

**Establishment-period climate (September-November)**

| Site | Year | Precipitation (mm) | Precipitation anomaly (mm) | Mean temperature (°C) | Temperature anomaly (°C) |
|---|---|---|---|---|---|
| Planthaven | 2021 | 137.4 | -114.6 | 15.0 | +1.1 |
| DDPSC FRS | 2022 | 205.5 | -44.7 | 13.7 | -0.3 |

**Reproductive-period climate (June-July)**

| Site | Year | Stand age | Precipitation (mm) | Precipitation anomaly (mm) | Mean temperature (°C) | Temperature anomaly (°C) | Mean maximum temperature (°C) | Maximum temperature anomaly (°C) |
|---|---|---|---|---|---|---|---|---|
| Planthaven | 2022 | 1 | 376.4 | +163.1 | 25.4 | +0.8 | 31.6 | +1.4 |
| Planthaven | 2023 | 2 | 243.8 | +30.5 | 24.3 | -0.3 | 31.0 | +0.8 |
| Planthaven | 2024 | 3 | 268.2 | +54.9 | 24.9 | +0.3 | 30.7 | +0.5 |
| DDPSC FRS | 2023 | 1 | 185.7 | -27.2 | 24.9 | +0.1 | 31.1 | +1.0 |
| DDPSC FRS | 2024 | 2 | 254.3 | +41.4 | 25.5 | +0.7 | 31.0 | +0.9 |
| DDPSC FRS | 2025 | 3 | 345.4 | +132.6 | 26.3 | +1.5 | 31.2 | +1.1 |

Climate data were obtained from PRISM.

#### Table S2. Effects of treatment and stand age on per-row IWG biomass, grain yield and Alfalfa biomass at Planthaven and DDPSC FRS.

**Type III fixed effects**

| Site | Response | Treatment | Stand age | Treatment × stand age |
|---|---|---|---|---|
| Planthaven | IWG biomass | **0.032** | **<0.001** | **0.032** |
| Planthaven | IWG grain yield | **0.017** | **0.003** | **0.009** |
| Planthaven | Alfalfa biomass | **0.001** | **< 0.001** | 0.609 |
| DDPSC FRS | IWG biomass | **<0.001** | **<0.001** | 0.051 |
| DDPSC FRS | IWG grain yield | **<0.001** | **<0.001** | 0.054 |
| DDPSC FRS | Alfalfa Biomass | **<0.001** | **<0.001** | 0.094 |

**Estimated marginal means for IWG**

| | | IWG biomass (g m⁻²) | | | IWG grain yield (g m⁻²) | | |
|---|---|---|---|---|---|---|---|
| Site | Stand age | IWG | IWG+N | IWG+Alf | IWG | IWG+N | IWG+Alf |
| Planthaven | 1 | 285.1a | 351.9a | 295.6a | 40.1a | 67.3a | 43.7a |
| Planthaven | 2 | 787.1a | 824.9a | 932.6a | 79.9a | 89.3a | 118.9a |
| Planthaven | 3 | 761.0ab | 491.7b | 1019.3a | 83.2ab | 36.6b | 133.0a |

| | | | | | | | |
|---|---|---|---|---|---|---|---|
| DDPSC FRS | 1 | 501.6b | 621.6ab | 797.8a | 85.1b | 78.1b | 154.1a |
| DDPSC FRS | 2 | 993.7b | 785.5b | 1332.0a | 37.7b | 31.6b | 79.7a |
| DDPSC FRS | 3 | 490.6b | 601.5ab | 703.0a | 12.1a | 14.9a | 28.9a |

**Estimated marginal means for Alfalfa biomass (g m$^{-2}$)**

| Site | Stand age | IWG+Alf | Alfalfa |
|---|---|---|---|
| Planthaven | 1 | 297a | 484a |
| Planthaven | 2 | 778a | 1020a |
| Planthaven | 3 | 39b | 394a |
| DDPSC FRS | 1 | 859a | 1024a |
| DDPSC FRS | 2 | 209b | 619a |
| DDPSC FRS | 3 | 152b | 446a |

Values are estimated marginal means. Within each site and stand age, means sharing a letter are not significantly different based on Sidak-adjusted pairwise comparisons ($P < 0.05$). The letter a denotes the highest statistical grouping; means may share letters when they are not significantly different. Per-row productivity is expressed per unit area of harvested IWG rows rather than per unit field area. IWG, intermediate wheatgrass; N, nitrogen; Alf, alfalfa.

#### Table S3. Effects of treatment and stand age on field-scale intermediate wheatgrass biomass, and grain yield, and alfalfa biomass at Planthaven and DDPSC FRS.

**Type III fixed effects**

| Site | Response | Treatment | Stand age | Treatment × stand age |
|---|---|---|---|---|
| Planthaven | IWG biomass | **<0.001** | **<0.001** | **0.042** |
| Planthaven | IWG grain yield | 0.133 | **0.006** | **0.014** |
| Planthaven | Alfalfa biomass | **<0.001** | **<0.001** | **<0.001** |
| DDPSC FRS | IWG biomass | **<0.001** | **<0.001** | 0.185 |
| DDPSC FRS | IWG grain yield | 0.551 | **<0.001** | 0.908 |
| DDPSC FRS | Alfalfa biomass | **<0.001** | **<0.001** | 0.057 |

**Estimated marginal means for IWG**

| | | IWG biomass (kg ha$^{-1}$) | | | IWG grain yield (kg ha$^{-1}$) | | |
|---|---|---|---|---|---|---|---|
| **Site** | **Stand age** | **IWG** | **IWG+N** | **IWG+Alf** | **IWG** | **IWG+N** | **IWG+Alf** |
| Planthaven | 1 | 2851a | 3519a | 1478a | 401ab | 673a | 219b |
| Planthaven | 2 | 7871a | 8249a | 4663b | 799a | 893a | 595a |
| Planthaven | 3 | 7610a | 4917b | 5096b | 832a | 366b | 665ab |
| DDPSC FRS | 1 | 5054ab | 6241a | 4028b | 851a | 781a | 956a |
| DDPSC FRS | 2 | 10123a | 8822ab | 6719b | 377a | 316a | 398a |
| DDPSC FRS | 3 | 4976ab | 6065a | 3531b | 121a | 149a | 145a |

**Estimated marginal means for Alfalfa biomass kg ha$^{-1}$**

| Site | Stand age | IWG+Alf | Alfalfa |
|---|---|---|---|
| Planthaven | 1 | 1468b | 4682a |
| Planthaven | 2 | 3694b | 9903a |
| Planthaven | 3 | 154b | 3885a |
| DDPSC FRS | 1 | 4296b | 10241a |
| DDPSC FRS | 2 | 1046b | 6187a |
| DDPSC FRS | 3 | 760b | 4456a |

Within each site and stand age, treatment means sharing a letter are not significantly different based on Sidak-adjusted pairwise comparisons (P < 0.05). Field-scale IWG biomass and grain yield account for the lower IWG row density in the IWG+Alf treatment. Alfalfa biomass comparisons include only IWG+Alf and alfalfa monoculture. IWG, intermediate wheatgrass; N, nitrogen; Alf, alfalfa.

#### Table S4. Effects of treatment on intermediate wheatgrass lodging at Planthaven and DDPSC FRS.

**Analysis of variance**

| Site and stand age | Effect | df | F-value | P-value |
|---|---|---|---|---|
| Planthaven - age 3 | Treatment | 2, 6 | 16.64 | **0.004** |
| DDPSC FRS - age 2 | Treatment | 2, 6 | 71.55 | **<0.001** |
| DDPSC FRS - age 3 | Treatment | 2, 24 | 64.55 | **<0.001** |
| DDPSC FRS - age 3 | Measurement | 2, 24 | 71.84 | **<0.001** |
| DDPSC FRS - age 3 | Treatment × measurement | 4, 24 | 16.27 | **<0.001** |

**Sidak-adjusted pairwise treatment comparisons**

| Site and stand age | Observation time | IWG vs. IWG+N | IWG vs. IWG+Alf | IWG+N vs. IWG+Alf |
|---|---|---|---|---|
| Planthaven - age 3 | Pre-harvest | **0.004** | 0.098 | 0.068 |
| DDPSC FRS - age 2 | Pre-harvest | **<0.001** | 0.997 | **<0.001** |
| DDPSC FRS - age 3 | Late May | 1.000 | 1.000 | 1.000 |
| DDPSC FRS - age 3 | Early July | **<0.001** | **<0.001** | **0.044** |
| DDPSC FRS - age 3 | Harvest | **<0.001** | **<0.001** | 0.261 |

Lodging was visually estimated as the percentage of plot area occupied by intermediate wheatgrass (IWG) stems positioned at less than 10 degrees from the soil surface. Values in section B are Sidak-adjusted P-values; bold values indicate P < 0.05.

#### Supplemental Table S5. Type III analysis of variance for root traits at Planthaven and DDPSC FRS.

| | | Planthaven | | | DDPSC FRS | | |
|---|---|---|---|---|---|---|---|
| **Trait** | **Effect** | **df** | **F** | **P** | **df** | **F** | **P** |
| **Total root length (cm)** | | | | | | | |
| | Treatment | 2, 6.00 | 2.86 | 0.134 | 2, 5.92 | 1.19 | 0.368 |
| | Depth | 2, 117.00 | 6.63 | **0.002** | 2, 116.53 | 50.94 | **<0.001** |
| | Position | 1, 117.00 | 2.31 | 0.131 | 1, 116.05 | 0.31 | 0.581 |
| | Treatment × Depth | 4, 117.00 | 0.59 | 0.674 | 4, 116.52 | 0.54 | 0.707 |
| | Treatment × Position | 2, 117.00 | 3.21 | **0.044** | 2, 116.05 | 0.29 | 0.752 |
| | Depth × Position | 2, 117.00 | 0.03 | 0.974 | 2, 116.05 | 0.35 | 0.708 |
| | Treatment × Depth × Position | 4, 117.00 | 0.83 | 0.508 | 4, 116.05 | 0.21 | 0.934 |
| **Root length density (cm/cm3)** | | | | | | | |
| | Treatment | 2, 6.00 | 2.45 | 0.167 | 2, 122.05 | 1.40 | 0.250 |
| | Depth | 2, 117.00 | 38.75 | **<0.001** | 2, 122.14 | 128.56 | **<0.001** |
| | Position | 1, 117.00 | 1.88 | 0.173 | 1, 122.03 | 0.16 | 0.687 |
| | Treatment × Depth | 4, 117.00 | 1.26 | 0.291 | 4, 122.14 | 0.48 | 0.754 |
| | Treatment × Position | 2, 117.00 | 2.30 | 0.105 | 2, 122.03 | 0.28 | 0.755 |
| | Depth × Position | 2, 117.00 | 0.24 | 0.790 | 2, 122.03 | 0.21 | 0.812 |
| | Treatment × Depth × Position | 4, 117.00 | 0.57 | 0.688 | 4, 122.03 | 0.23 | 0.921 |
| **Specific root length (cm/g)** | | | | | | | |
| | Treatment | 2, 124.00 | 0.10 | 0.904 | 2, 5.98 | 0.31 | 0.744 |
| | Depth | 2, 124.00 | 3.09 | **0.049** | 2, 116.38 | 0.71 | 0.495 |
| | Position | 1, 124.00 | 2.16 | 0.144 | 1, 115.90 | 0.09 | 0.759 |
| | Treatment × Depth | 4, 124.00 | 0.98 | 0.422 | 4, 116.37 | 0.24 | 0.915 |
| | Treatment × Position | 2, 124.00 | 1.13 | 0.327 | 2, 115.90 | 0.15 | 0.863 |
| | Depth × Position | 2, 124.00 | 1.05 | 0.354 | 2, 115.90 | 3.31 | **0.040** |
| | Treatment × Depth × Position | 4, 124.00 | 0.52 | 0.722 | 4, 115.90 | 0.80 | 0.529 |
| **Average root diameter (mm)** | | | | | | | |
| | Treatment | 2, 6.00 | 0.15 | 0.862 | 2, 5.94 | 1.40 | 0.319 |
| | Depth | 2, 117.00 | 4.96 | **0.009** | 2, 116.35 | 5.12 | **0.007** |
| | Position | 1, 117.00 | 2.57 | 0.112 | 1, 115.86 | 0.21 | 0.644 |
| | Treatment × Depth | 4, 117.00 | 3.36 | **0.012** | 4, 116.33 | 0.52 | 0.719 |
| | Treatment × Position | 2, 117.00 | 2.13 | 0.124 | 2, 115.86 | 0.23 | 0.795 |
| | Depth × Position | 2, 117.00 | 13.95 | **<0.001** | 2, 115.86 | 3.89 | **0.023** |
| | Treatment × Depth × Position | 4, 117.00 | 1.09 | 0.367 | 4, 115.86 | 0.29 | 0.883 |
| **Root mass (g)** | | | | | | | |
| | Treatment | 2, 124.00 | 0.90 | 0.408 | 2, 125.00 | 0.31 | 0.737 |
| | Depth | 2, 124.00 | 26.49 | **<0.001** | 2, 125.00 | 10.84 | **<0.001** |
| | Position | 1, 124.00 | 27.25 | **<0.001** | 1, 125.00 | 1.66 | 0.200 |
| | Treatment × Depth | 4, 124.00 | 0.36 | 0.837 | 4, 125.00 | 0.55 | 0.697 |
| | Treatment × Position | 2, 124.00 | 1.78 | 0.172 | 2, 125.00 | 0.22 | 0.803 |
| | Depth × Position | 2, 124.00 | 20.40 | **<0.001** | 2, 125.00 | 2.14 | 0.122 |
| | Treatment × Depth × Position | 4, 124.00 | 0.33 | 0.860 | 4, 125.00 | 0.51 | 0.725 |

Separate linear mixed-effects models were fitted for each root trait and site. Fixed effects included treatment, soil depth, sampling position, and their interactions, with block/plot as random effects. df = numerator and denominator degrees of freedom. Bold P-values indicate significant effects ($P < 0.05$). P-values below 0.001 are displayed as <0.001.
